# RELIEF: a structured multivariate approach for removal of latent inter-scanner effects

**DOI:** 10.1101/2022.08.01.502396

**Authors:** Rongqian Zhang, Lindsay D. Oliver, Aristotle N. Voineskos, Jun Young Park

## Abstract

Combining data collected from multiple study sites is becoming common and is advantageous to researchers to increase the generalizability and replicability of scientific discoveries. However, at the same time, unwanted *inter-scanner biases* are commonly observed across neuroimaging data collected from multiple study sites or scanners, rendering difficulties in integrating such data to obtain reliable findings. While several methods for handling such unwanted variations have been proposed, most of them use univariate approaches that could be too simple to capture all sources of scanner-specific variations. To address these challenges, we propose a novel multivariate harmonization method, called RELIEF (**RE**moval of **L**atent **I**nter-scanner **E**ffects through **F**actorization) for estimating and removing both explicit and latent scanner effects. Our method is the first approach to introduce the simultaneous dimension reduction and factorization of interlinked matrices to a data harmonization context, which provides a new direction in methodological research for correcting inter-scanner biases. Analyzing diffusion tensor imaging (DTI) data from the Social Processes Initiative in Neurobiology of the Schizophrenia (SPINS) study and conducting extensive simulation studies, we show that RELIEF outperforms existing harmonization methods in mitigating inter-scanner biases and retaining biological associations of interest to increase statistical power. RELIEF is publicly available as an R package.

## 1. Introduction

It is increasingly common in neuroimaging and genomics to combine data collected from multiple study sites to increase the power and the reproducibility of scientific discoveries. However, combining such data comes with unwanted non-biological variations that need to be removed for successful data integration. In neuroimaging, this is often characterized by inter-scanner biases (scanner effects) when subject data are obtained by using different magnetic resonance imaging (MRI) scanners with different optimization protocols. These inter-scanner biases have been shown to be present in most neuroimaging data types, including diffusion [1, 2], structural [3, 4], and functional [5] MRI. These terms are analogous to *batch effects* in genomic studies that are observed with genome-wide microarray or RNA sequencing data with different sample preparation and sequencing methods.

There have been numerous efforts in statistics, such as ComBat, to capture and remove these unwanted variations while increasing the signal-to-noise ratio [6, 7, 8, 9, 10]. ComBat [6] is a popular regression-based batch correction approach first motivated from microarray data, and has been promising in removing inter-scanner biases in many neuroimaging data types, including fractional anisotropy and mean diffusivity [8], cortical thickness [9], and functional connectivity [10]. In ComBat, scanner effects are characterized by an additive scanner effect (location) and a multiplicative scanner effect (scale) for each feature. While a regression model is used in each feature, ComBat uses empirical Bayes to stabilize estimates across features and provides robustness in the case of small within-scanner sample sizes [6]. In addition to showing its utility in various neuroimaging data types, ComBat has been extended to harmonize imaging data collected in a longitudinal manner [11], to preserve non-linear age trajectories of cortical thickness data in mega analysis in cross-sectional studies [12]. It is also a versatile method that allows for harmonization even without the need to share original data from a study site with other sites, which relaxes concerns about data privacy [13].

The ComBat’s location-scale model is simple and interpretable, but, from the statistical perspective, it is insufficient to capture all sources of scanner effects. The heterogeneity in *covariances* across different sites or scanners has been overlooked in the neuroimaging literature, and such heterogeneity might also lead to decreased statistical power. ComBat is oversimplified by the assumption that additive scanner effects can be explained by only an intercept for each scanner and feature. Recently, a new harmonization method called CovBat [14] was proposed to address covariance heterogeneity in multi-site, multi-scanner studies by extending ComBat. It applies ComBat twice: first to the original data, then to the principal component scores from the residual matrix. CovBat is an important development that expanded the scope of statistical harmonization to address heterogeneous covariances, and it has been shown to be more efficient than ComBat, as expected [14, 15]. However, CovBat implicitly assumes that the covariance scanner effect is contained within the eigenspace of the residual matrix only, in the form of a location-scale model. As Chen et al. [14] noted, this assumption may limit the ability of CovBat to characterize all sources of covariance heterogeneity, which we also show in this paper.

The method for harmonizing covariances across scanners can be understood using the latent variable formulation [14]. Singular value decomposition (SVD) or principal component analysis (PCA) are commonly used techniques for removing or adjusting for non-biological variations not explicitly specified by scanner information. SVA (Surrogate Variable Analysis) is a method that was originally developed for genomic studies [16] and then adapted to neuroimaging studies [17]. SVA includes latent factors of unwanted variation as surrogate variables, which are not associated with the biological covariates of interest. Instead of using explicit variables to denote scanner effects, SVA identifies and estimates scanner or other non-biological artifacts through permutation testing, then removes them as surrogate variables. RAVEL [17] is a scanner effect correction method for neuroimaging data inspired by RUV (remove unwanted variables) [18]. RAVEL estimates and removes unwanted variation factors by using negative controls, which are features (e.g. genes, voxels, etc.) that are known *a priori* to be unassociated with the variables of interest [8, 17]. This method applies SVD to obtain latent factors of unwanted variations in the control regions and then removes the latent factors and corresponding effects in the test regions. These approaches that apply low-rank factorization methods to all study subjects’ imaging features are fundamentally limited to addressing *scanner-specific* latent effects. At the same time, efforts to identify low-rank factors for study subjects from the same scanner may overkill biological variations.

In this paper, we propose a novel harmonization method, called RELIEF (**RE**moval of **L**atent **I**nter-scanner **E**ffects through **F**actorization) to distinguish loadings shared across scanners (which should be preserved) from loadings specific to scanners (which should be removed), which enhances the current understanding of inter-scanner biases. We formulate latent scanner effects from the perspective of linked matrix factorization by extending the recent work of Park and Lock [19] in the harmonization context. It aligns with growing methodological developments on simultaneous dimension reduction and factorization of multi-modal data (e.g., [20, 21, 22]), which has also been shown to be promising in neuroimaging data [23]. Through extensive data analyses and simulations, we show our proposed method has superior performance in identifying and removing latent unwanted variations specific to each scanner, thus leading to covariance homogeneity across scanners and increasing statistical power compared to existing methods. Also, our estimation procedure is scalable and takes only a few seconds to implement, which supports its practical utility.

The rest of the paper is organized as follows. Section 2 describes our proposed method, RELIEF, and compares it to existing harmonization methods. In Section 3, we apply our method to the fractional anisotropy (FA) and mean diffusivity (MD) data from the Social Processes Initiative in the Neurobiology of the Schizophrenia(s) (SPINS) study, where study subjects were collected from multiple sites and scanners. We compare RELIEF to other harmonization methods using a comprehensive evaluation framework. Section 4 conducts extensive simulations to evaluate performances in terms of Type 1 error rate and statistical power. We conclude with some points of discussion in Section 5.

## 2. Methods

### 2.1. Notation and setup

We let *i* = 1, …, *M* denote the index for each scanner (batch), *j* = 1, … *n*_*i*_ denote the subject index in *i*th scanner 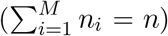, and *v* = 1, …, *V* denote the index for imaging features. We let **x**_*ij*_ be the *q*-dimensional covariate vector for *j*th subject in *i*th scanner (e.g. age and sex). *y*_*ijv*_ is the *v*th imaging feature of the *j*th subject of the *i*th scanner. By stacking all observations of **x**_*ij*_, we let **X** be a *n × q* matrix of *q* covariates observed for *n* study subjects. Similarly, let **Y** be a *V × n* data matrix of *V* features. Then, to group the subjects from the same scanner together, we consider *{***Y**_*i*_ : *V × n*_*i*_|*i* = 1, …, *M}* a partition of **Y**. The matrices can be concatenated to form a matrix **Y** = [**Y**_1_; **Y**_2_; … ; **Y**_*M*_]. We will use this notation for a general *V × n* matrix throughout this paper.

### 2.2. Existing harmonization methods

#### 2.2.1. Adjusted residuals (AdjRes)

The simplest approach to model inter-scanner bias is to use a regression-based approach to characterize additive scanner-specific deviations for each feature. AdjRes considers the following specifications,

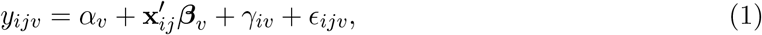

where, for the *v*th feature, *α*_*v*_ is the intercept, ***β***_*v*_ is the regression coefficients for **x**_*ij*_, and *ϵ*_*ijv*_ is a Gaussian noise. The parameters *α*_*v*_, ***β***_*v*_, *γ*_*iv*_ can be estimated by the least squares method. The scanner-specific means, *γ*_*iv*_ needs to be removed and the harmonized data is constructed by 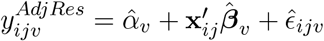.

#### 2.2.2. ComBat

ComBat seeks to remove the additive and multiplicative scanner effects [6]. For the *v*th feature, ComBat characterizes the additive and multiplicative scanner effects by

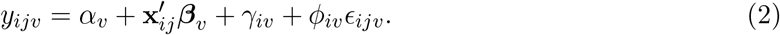

In Equation (2), the scanner effects are characterized by *γ*_*iv*_ (the additive scanner effect) and *ϕ*_*iv*_ (the multiplicative scanner effect). After obtaining 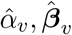 via least squares, ComBat estimates scanner effects in locations (i.e., 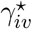) and scales (i.e., 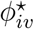) via empirical Bayes for each feature separately, providing stable and robust estimations of these parameters in the case of small within-scanner sample sizes [6]. The ComBat-harmonized data is defined by 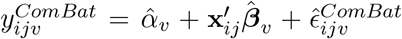, where

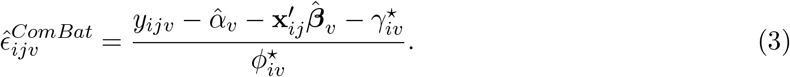

#### 2.2.3. CovBat

ComBat is fundamentally a univariate method that only removes scanner effects in mean and variance and it does not address the heterogeneity in covariances. To further improve data quality by addressing covariance heterogeneity, Chen et al. [14] recently proposed a multivariate batchcorrection method extending ComBat, called CovBat, which takes the covariance of multiple imaging features into consideration for harmonization of covariances.

Inspired by how ComBat mitigates the difference between the variance within each scanner and the pooled variance, CovBat shifts the within-scanner covariance to the pooled covariance by using principal component (PC) and PC scores. CovBat’s harmonization procedure is summarized as follows. First, ComBat is applied to full imaging data, yielding ComBat-residuals as in Equation (3) with homogeneous variances across scanners. CovBat then conducts the eigendecomposition on the sample covariance of Combat-residuals and applies ComBat again to each eigenvector to remove heterogeneous means and variances, which yields CovBat-residuals with an additional source of scanner effect removed. The final CovBat-harmonized data is 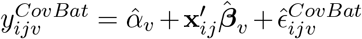. Cov-Bat assumes that the covariance scanner effects can be captured by the location-scale adjustments to the principal components of the residuals. Despite its efficiency, we point out that CovBat’s assumption might not be sufficient to characterize all sources of covariance heterogeneity.

### 2.3. New method: RELIEF(REemoval of Latent Inter-scanner Effects through Factorization)

We first characterize three sources of scanner effects (additive mean (location), additive latent, and multiplicative scanner effects (scale)) via an additive multivariate model illustrated in Figure We assume that the data matrix **Y** consists of

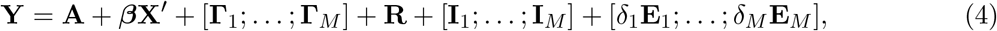

where **A** is the intercept matrix (rank of 1), ***β*** is a *V × q* matrix of regression coefficients (rank of min(*V, q*)), and [**Γ**_1_; … ; **Γ**_*M*_] is a matrix of additive scanner effects (locations) for each feature (rank of *M*), where elements of each row of **Γ**_*i*_ takes the same value. Note that **A** + ***β*X**^*′*^ + [**Γ**_1_; … ; **Γ**_*M*_] in Equation (4) corresponds to the collection of 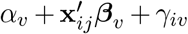 in Equation (1) across all imaging features. The RELIEF model assumes that *ϵ*_*ijv*_ in Equation (1) is decomposed into three additive variations. Specifically:

- **R** is a *V ×n* matrix of the latent structure explaining shared variations across all scanners but not explained by covariate effects. It includes (i) non-linear covariate effects from **X** or (ii) any additional variations due to unobserved covariates. From the viewpoint of scanner-effect correction, this should be preserved after harmonization.
- **I**_*i*_ is a *V × n*_*i*_ matrix of latent variations explaining latent scanner effects in the *i* scanner beyond scanner-specific means **Γ**_*i*_ (locations). This might include any non-linear scanner effects [24] or any contributions. It should be removed after harmonization.
- *δ*_*i*_**E**_*i*_ is a *V × n*_*i*_ noise matrix, and each element of **E**_*i*_ is assumed have a unit variance. *δ*_*i*_ characterizes the variance heterogeneity as specified in ComBat, which has shown to be promising in neuroimaging. From the viewpoint of scanner-effect correction, *δ*_*i*_s should be standardized to have a common variance across scanners.

Throughout this paper, we assume **R** as well as each of **I**_1_, …, **I**_*M*_ to be low rank, and estimate their ranks using a model-based approach.

Our approach is summarized by (i) removing scanner, feature-specific means and obtaining covariate effects first, (ii) standardizing the data matrix to have homogeneous variance, (iii) decomposing it into scanner-specific and scanner-independent factors, and (iv) reconstructing harmonized data.

Steps (i) and (ii) are achieved through the preprocessing step. We obtain least square estimates 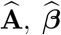 and 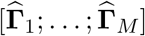 by using the two-step regression. Specifically, we first fit GLM using the intercept and covariates (**A** and ***β***) and obtain residuals 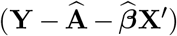. Then, using the residuals from the first step, we remove scanner-specific means for each feature (**Γ**)to obtain the second-step residuals 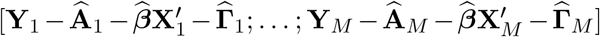 to be used in subsequent steps. When the variability of the second-step residuals differs across features, we can easily scale each residual by its residual standard deviation, apply steps (iii) and (iv), and scale back each feature.

Step (iii) is achieved by simultaneous dimension reduction and factorization methods proposed for cancer genomics in Park and Lock [19] and Lock et al. [22]. We first scale each residual matrix from the last step by 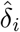 in order to make the residual variances homogeneous across *i* = 1, …, *M* : 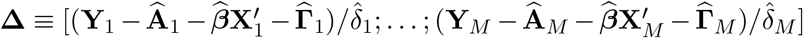. Following Park and Lock [19] and Lock et al. [22], we estimate 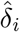 by the median of the singular values of residual matrices for each scanner divided by the square root of the median of the Marcenko-Pastur distribution [25]. Provided that 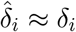, we first note that **Δ** is represented by

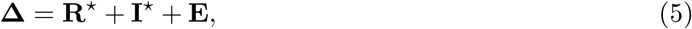

where 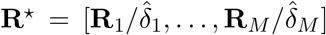 is a variation shared across all scanners, 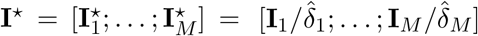 are individual variations shared only in each scanner.

From model (5), 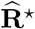 and 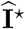 are obtained by

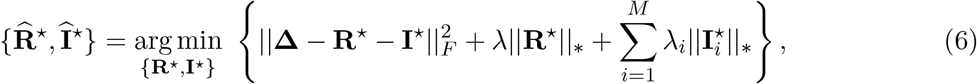

where 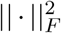 and || · ||_*_ are the squared Frobenious norm (sum of squared elements) and the nuclear norm (sum of singular values), respectively. The nuclear norm penalties in Equation (6) ensure that the resulting estimates 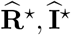 are low-rank [26]. Although tuning *λ* and *λ*_*i*_s may be tricky, we use the recommended values from Park and Lock [19] by setting 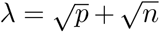 and 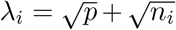, which was shown to perform well with independent Gaussian noise. With *λ* and *λ*_*i*_s specified, an iterative algorithm can be applied to estimate **R**^⋆^ and 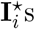.

In Step (iv), we scale 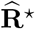 back to 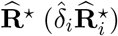 to make sure 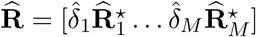 is in the original scale. To keep the noise variance homogeneous, we scale 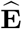 to 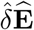, where 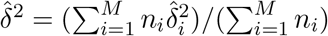 is the weighted mean of scanner-specified noise variance. Therefore, the final harmonized data is given by intercepts

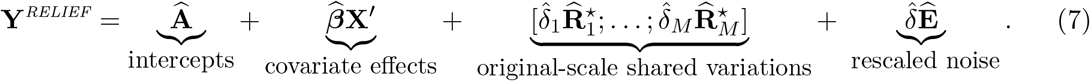

### 2.4. Using covariates in RELIEF

When a primary interest is to test for an association with a covariate of interest, including the covariate in RELIEF may lead to an inflated false positive rate. It is because our objective function (6) does not enforce scores of 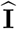 to be independent of the covariate of interest. Therefore, we suggest not including variables of interest when applying RELIEF. In practice, we found that even not including any covariates does not result in a noticeable difference. It is because the covariate effects are actually low-rank (with rank equal to the number of covariates) scanner-independent patterns and are captured by **R** (in high signal-to-noise ratio (SNR)) or by **E** (in low SNR), provided that covariates are independent to scanner information. However, we note that ComBat does not suffer from potential inflated false positives if the variables of interest and the indicator variable for additive scanner effect are linearly independent [27].

### 2.5. Preventing distorted covariate effects in RELIEF

Many existing harmonization methods, including AdjRes, ComBat, CovBat, and RELIEF, account for explicit covariate effects in the form of regression, but there might be hidden covariate effects from unobserved covariates. For downstream analyses, it is critical to preserve these effects in the original scale. In RELIEF, such effects correspond to the **R** term, and therefore, we scale 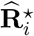 back to 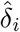 in Equation (7) although *δ*_*i*_ were used to characterize variance heterogeneity.

We point out that ComBat (and CovBat that uses ComBat in the first step) models *observed* covariate effects only and all *unobserved* covariate effects are attributed to the residuals. Since residuals are eventually scaled differently for each scanner/site in the harmonization steps, ComBat and CovBat could be prone to distorted covariate effects for unobserved covariates after harmonization, especially when variance heterogeneity across scanners is evident.

## 3. Data Analysis

### 3.1. Data preparation and preprocessing

We used diffusion tensor imaging (DTI) data from Social Processes Initiative in the Neurobiology of the Schizophrenia(s) (SPINS) study to empirically evaluate RELIEF’s performance. The study subjects consisted of 256 individuals with schizophrenia spectrum disorders (SSDs) and 175 controls. Subjects were 18–55 years old, and 268 of the participants were males (163 females). Participants with SSDs met DSM-5 diagnostic criteria for schizophrenia, schizoaffective disorder, schizophreniform disorder, delusional disorder, or psychotic disorder not otherwise specified, assessed using the Structured Clinical Interview for DSM (SCID-IV-TR), and had no change in antipsychotic medication or decrement in functioning/support level in the 30 days prior to enrollment. Controls did not have a current or past Axis I psychiatric disorder, excepting adjustment disorder, phobic disorder, and past major depressive disorder (over two years prior; presently unmedicated), or a first-degree relative with a history of psychotic mental disorder. Additional exclusion criteria included a history of head trauma resulting in unconsciousness, a substance use disorder (confirmed by urine toxicology screening), intellectual disability, debilitating or unstable medical illness, or other neurological diseases. Participants also had normal or corrected-to-normal vision. All participants signed an informed consent agreement and the protocol was approved by the respective research ethics and institutional review boards. All research was conducted in accordance with the Declaration of Helsinki.

The scans were acquired at three different imaging sites, including the Centre for Addiction and Mental Health (CAMH), Maryland Psychiatric Research Center (MPRC), and Zucker Hillside Hospital (ZHH). General Electric 3T MRI scanners were used at CAMH and ZHH (750w Discovery and Signa, respectively), and the Siemens Tim Trio 3T MRI scanner at MPRC. However, during the middle of the study, all study sites switched to Siemens Prisma 3T scanners for data collection. A high-angular resolution axial EPI dual spin echo sequence diffusion scan was acquired on all scanners. Within the limits of scanner hardware, parameters were prospectively harmonized as follows: 60 gradient directions, b=1000, 5 b=0 images, TR=8800 ms (one scanner TR=17000 ms), TE=85 ms, FOV=256 mm; in-plane matrix 128×128, and 2.0 mm isotropic voxels. All images were preprocessed using the same pipeline across sites. Skull-stripping was performed via a two-step process combining FSL (BET) and AFNI to optimize brain extraction, after which MRtrix3 (dwi2mask) was used for brain masking. FSL eddy was used for eddy current-induced distortion and motion correction, including volume-to-volume and within-volume movement [28]. Eddy models the effects of participant movement and diffusion eddy currents simultaneously, predicting undistorted data using a Gaussian Process. Eddy also outputs quality control metrics, including average absolute motion (mm) for each participant as one measure of volume-to-volume movement. Fieldmap-free susceptibility distortion correction was performed using BrainSuite (BDP; [29]). Outputs were visually inspected after each preprocessing step to ensure data quality.

Participants’ white matter tracts were reconstructed using deterministic unscented Kalman Filter (UKF) tractography [30] in 3D Slicer (https://github.com/SlicerDMRI). The ORG (O’Donnell Research Group) white matter atlas [31] was used to parcellate fibers into anatomical tracts. This atlas has been validated across different scanners and protocols (e.g., number of gradient directions, spatial resolutions, b-values;[32]). Metrics were included from 56 deep white matter fiber tracts from the association, cerebellar, commissural and projection tracts (the cortico-ponto-cerebellar tract was excluded due to parcellation issues), and 16 superficial tract categories according to the brain lobes they connect, resulting in *V* = 72 features. Mean FA values and mean diffusivity (MD) values were calculated along each tract. FA measures the degree to which diffusion of water molecules is restricted by microstructural elements such as cell bodies, axons, myelin, and other constituents of cytoskeleton [33]. MD is a measure of the magnitude of water diffusion, independent of direction [34]. Visual quality control was performed after initial tractography, registration to the ORG atlas, and tract creation. Data from seven participants were excluded on the basis of missing or poor tractography for *>* 15 tracts across the whole brain.

Since the number of samples from Siemens Tim Trio are small, we used images from two scanner types (GE and SP) in our analysis. Participants without DTI data were also excluded from the study. The final sample consists of 351 subjects across 2 scanner types, with 172 subjects imaged on scanners manufactured by GE (67 females, 111 patients, age 18-55), 179 on Prisma scanners manufactured by Siemens (71 females, 98 patients, age 18-55).

### 3.2. Results

We harmonized data by using RELIEF, ComBat, CovBat, and AdjRes. We used age, age^2^, gender, diagnosis, an interaction between age and gender (age*×*gender) and an interaction between age and diagnosis (age*×*diagnosis) to model covariate effects in harmonization.

Figure 2 shows the heatmap of the estimated latent scanner effects 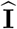 of RELIEF for the FA and MD data from the SPINS study. As RELIEF’s crucial components, the latent scanner effects are identified and removed to reduce the inter-scanner variations directly. In Figure 2, the most scanner-specific variations were attributed to Siemens Prisma for both FA and MD. To investigate the potential sources of latent Siemens Prisma-specific variations in relation to existing non-biological information, we applied hierarchical clustering to the site subgroups of 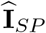 in Figure 2 and reordered subjects within Siemens Prisma so that 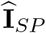 within the same site were arranged together. We observed the latent scanner effects within each site tended to share similar patterns, which suggests that the variations in 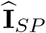 are highly associated with sites.

**Figure 1:**
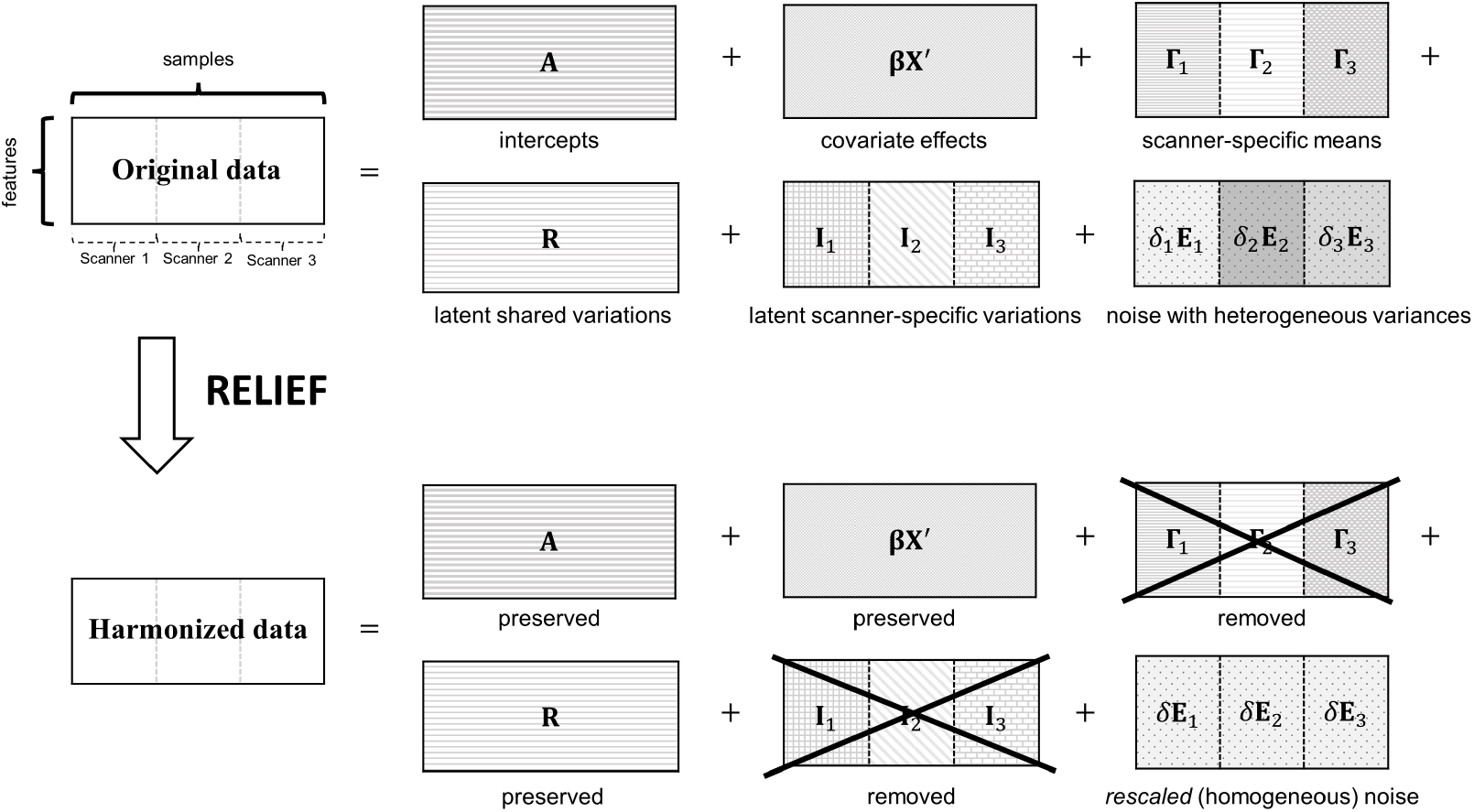
Overview of RELIEF using data consisting of three scanners for illustrations. It decomposes original data as (i) covariate effects, (ii) scanner-specific means (locations), (iii) latent shared variations, (iv) latent scanner-specific variations, and (v) noise with heterogeneous variances (scales). For harmonization purposes, RELIEF removes (ii) and (iv) specific to scanners and homogenizes (v).

**Figure 2:**
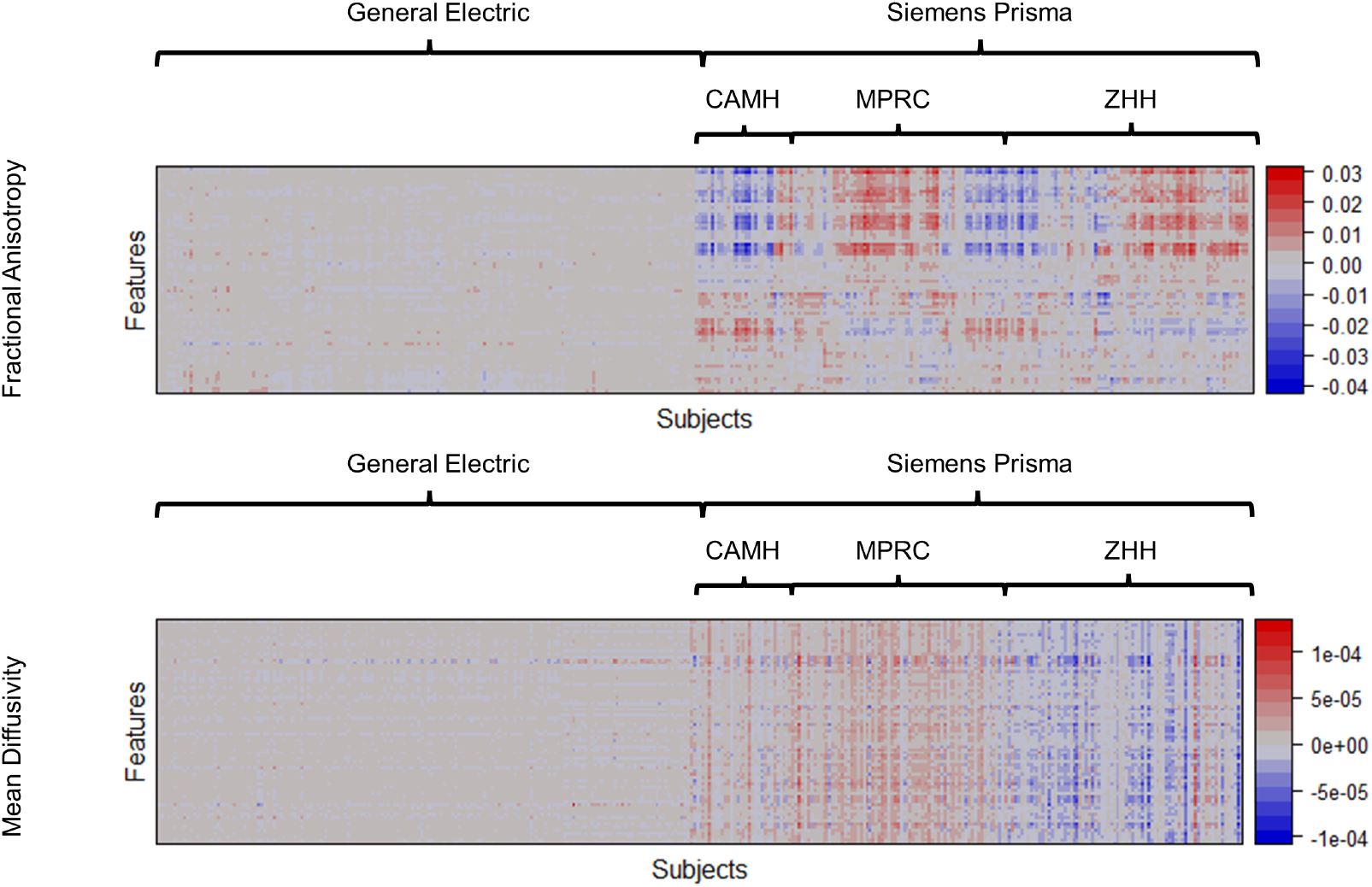
Heatmaps of the estimated latent scanner-specific variations 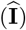 of the FA and MD from the SPINS study. For visualizations, imaging features were reordered by applying hierarchical clustering; subjects scanned by General Electric 3T were reordered separately, and subjects scanned by Siemens Prisma 3T were reordered within each site subgroup (CAMH, MPRC, and ZHH). Feature indices were also reordered by applying hierarchical clustering to concatenated **I** for FA and MD. RELIEF identified substantial variations present mostly on Siemens Prisma but not on General Electric, and the variations are highly associated with sites.

We performed statistical analysis to quantify the relationship between existing non-biological information, including site information and motion parameters. In Figure 3 (a) and (b), we performed one-way ANOVA to compare different latent scanner effects of the Siemens Prisma across sites for FA and MD data respectively. We found that the latent factors of most features specific to Simens Prisma were highly associated with the sites, particularly for MD data. In Figure 3 (c) and (d), we performed correlation tests between the latent scanner effects in the Siemens Prisma scanner and the motion parameter for FA and MD data, respectively. We calculated the average absolute motion from the reference volume (in mm) to represent subject motion during the scan and averaged it for the six motion parameters (three translations and three rotations). Our findings revealed that the latent factors showed no significant associations with the motion parameter.

**Figure 3:**
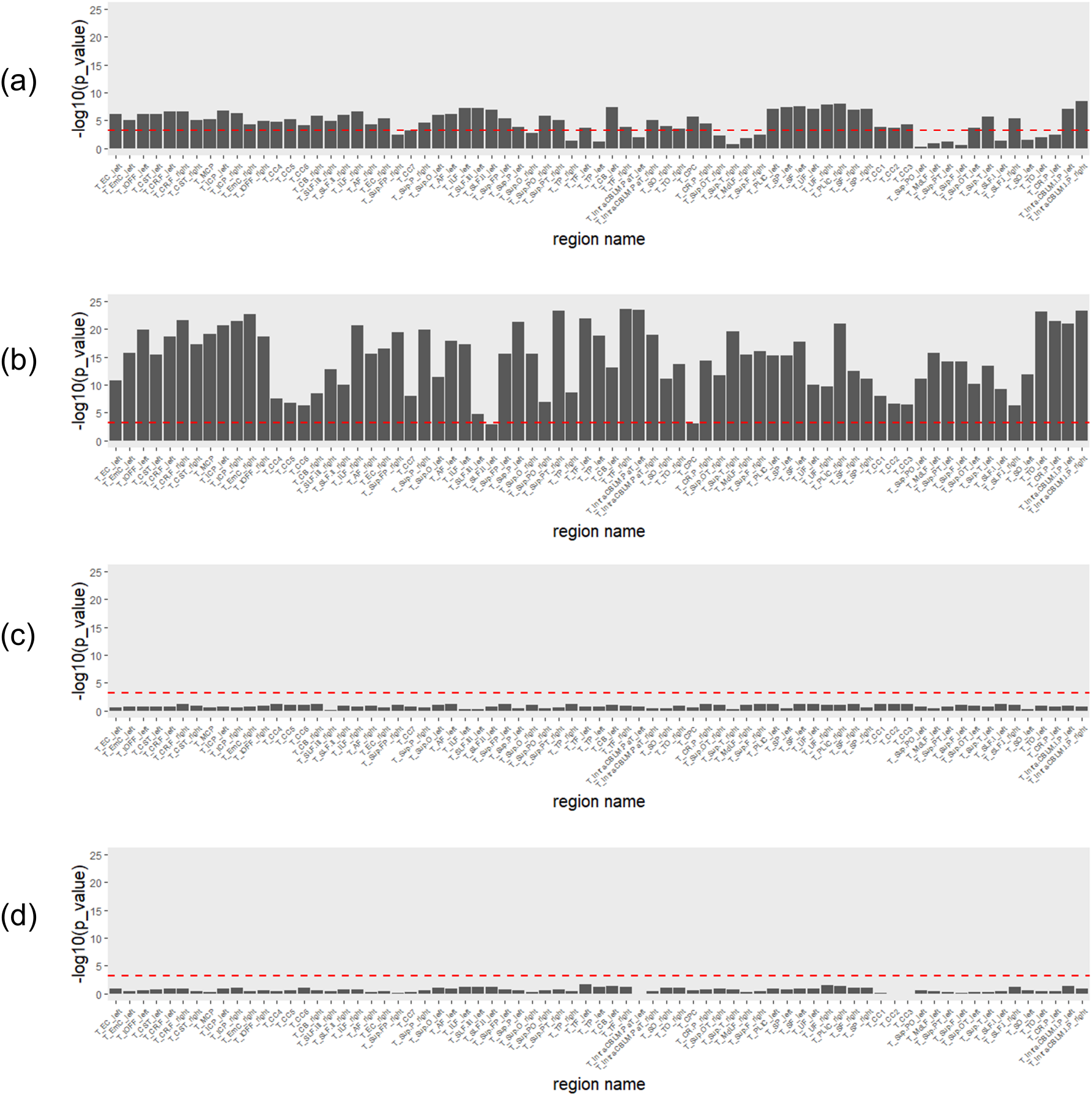
Investigating the potential sources of latent Siemens Prisma-specific variations in relation to existing non-biological information (site information and a motion parameter). (a) and (b) show one-way ANOVA *p* values of (a) FA and (b) MD in relations to 3 study sites. (c) and (d) show *p* values for the correlation between latent factors and the average absolute motion from the reference volume (in mm) for (c) FA and (d) MD. All *p* values were negative log-transformed (with base 10) for visualizations. The red dashed horizontal line is Bonferroni-corrected threshold (0.05*/*72 ≈ 6.9 *×* 10^−4^). The region names agree with the features’ order in Figure 2.

Overall, our analyses provided insights into how existing non-biological information can impact the interpretation of latent scanner-specific variations.

To visualize whether most variations in the data are still associated scanner after harmonization, we applied two unsupervised data reduction techniques: principal component analysis (PCA) and t-distributed stochastic neighbor embedding (t-SNE) to the original and harmonized FA and MD data from diffusion tensor imaging (DTI). As a nonlinear technique, t-SNE emphasizes preserving the variations in the local structure of the data, while PCA focuses more on preserving variations in the overall data set. The data projected into the first two PCs/dimensions are presented in Figure

For raw data, we observed that most variations are clearly explained by the scanner information (General Electric vs. Siemens Prisma). For AdjRes and ComBat, despite evidence of higher data quality, there is heteroscedasticity of ellipses across scanners, which indicates that there are still unremoved latent scanner effects. For CovBat and RELIEF, both PC scores and t-SNE scores appear to be distributed similarly across scanners, which suggests the variations associated with scanners are substantially removed.

To evaluate if scanner-specific latent patterns are well-removed, we computed the empirical covariances by scanners as well as the difference between two scanner-specific covariances. Figure 5 shows that the covariance differences remain notable in AdjRes harmonized data. ComBat and CovBat performed slightly better than AdjRes in mitigating covariance scanner effects. Notably, however, these covariance differences are considerably reduced with RELIEF. We also quantified these differences in covariances by the Frobenius norm of the scanner-specific covariance matrices. For FA, the norm for RELIEF was the lowest (**3.70**) followed by CovBat (5.77), ComBat (6.19) and AdjRes (8.10). For MD, the norm for RELIEF was also the lowest (**1.45**) followed by CovBat (2.29), ComBat (4.15) and AdjRes (8.74). These results suggest the superior performance of RELIEF in constructing homogeneous covariances.

**Figure 4:**
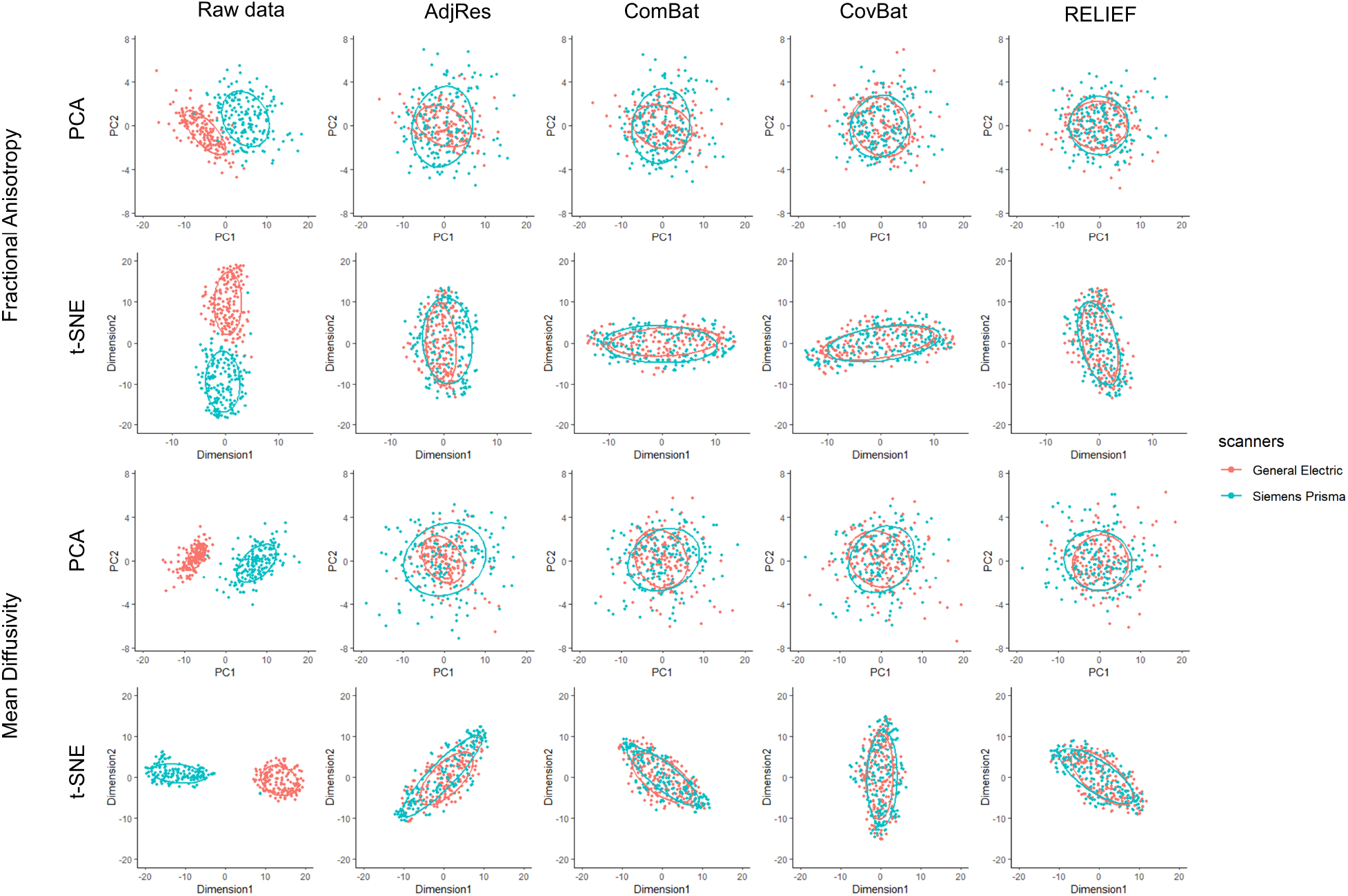
Scatterplots of principal component scores and t-SNE scores before and after applying harmonization to the SPINS DTI data.

**Figure 5:**
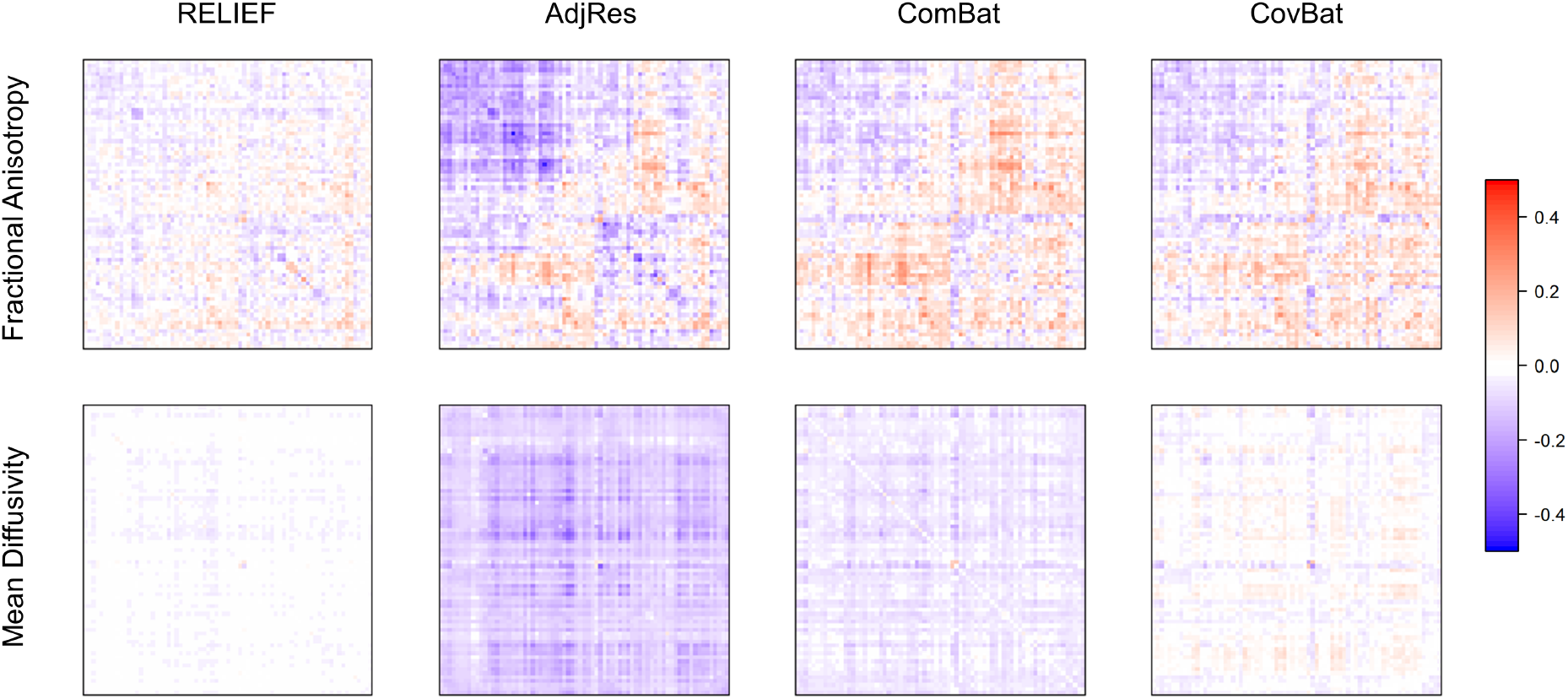
The difference of scanner-specific covariance matrices for harmonized SPINS data (GE−SP). The order of the features agrees with Figure 2 and 3. The x-axis and y-axis indicate regions of interest, which are explicitly illustrated in the x-axis of Figure 3. The color bar shows the range of values of the differences in covariances. RELIEF reveals the lowest difference between two covariances.

We also used Quadratic Discriminant Analysis (QDA) to evaluate how data harmonized using each approach predicts scanners. A harmonization method that performs *better* in removing scanner effects will result in *worse* predictive performance. Using machine learning methods to predict scanners from harmonized data has been adopted in previous work in evaluating the performance of different harmonization methods [9, 14]. We chose QDA because the classifier is constructed based on the mean vectors and covariance matrices only, where differences in predictive performances are attributed to the harmonization of scanner-specific means and covariances. Using leave-one-out cross-validation, we computed the average accuracy, ROC curve and its area under the curve (AUC) for each harmonized data after regressing out covariate effects. For FA, the RELIEF method achieved the lowest prediction accuracy (**49.6%**) close to a random prediction, followed by Cov-Bat (59.3%), ComBat (66.1%) and AdjRes (70.1%). For MD, RELIEF also achieved the lowest prediction accuracy (**61.0%**) followed by CovBat (82.6%), ComBat (83.2%) and AdjRes (87.5%) The results of the AUC, shown in Figure 6, were similar to the prediction accuracy, suggesting the lowest AUC for RELIEF.

**Figure 6:**
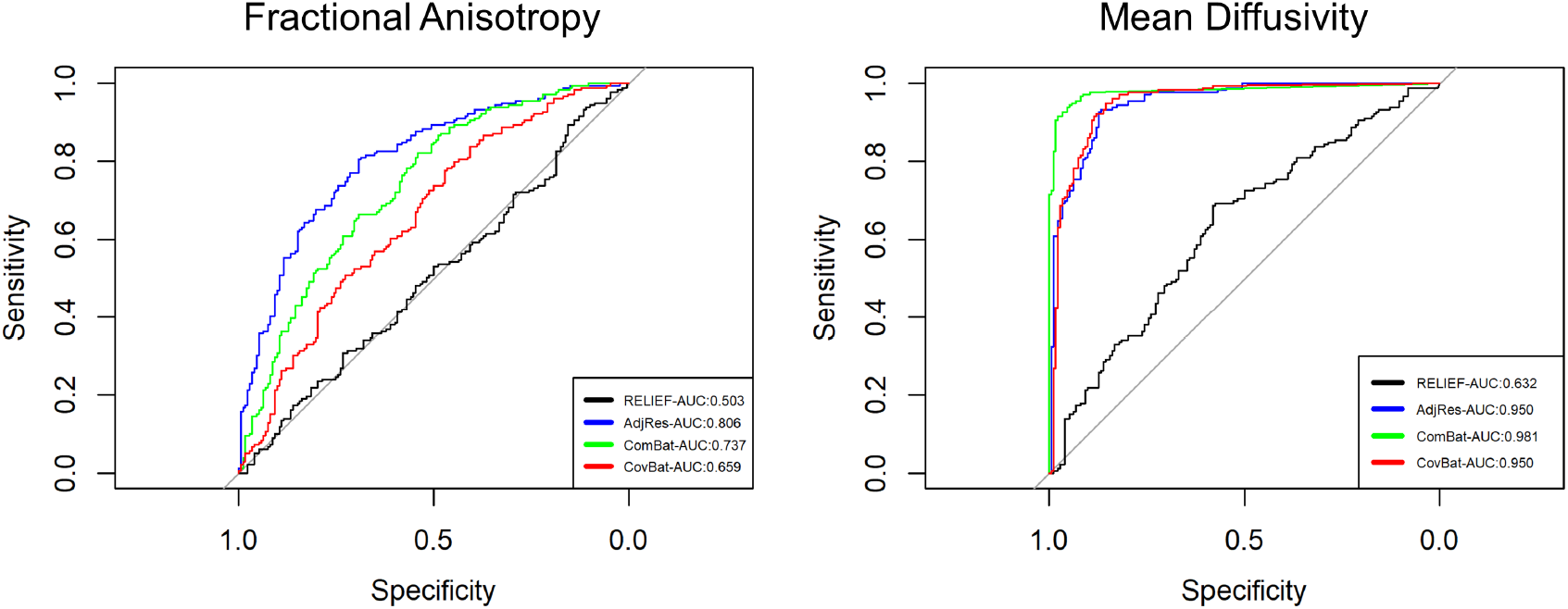
The ROC curves for predicting scanners by using SPINS data harmonized by different methods. We used QDA as a classifier and leave-one-out cross-validation (LOOCV) to obtain individualized predictions. The ROC curve of the RELIEF was closest to the diagonal line, suggesting that it successfully harmonized latent inter-scanner biases.

Lastly, we investigated whether RELIEF preserves the biological variability in the data. This step is necessary because the multivariate harmonization methods could be prone to potentially overkilling too much variation, including biological variations. Here, we evaluated whether the different harmonization methods maintain the biological associations of interest through multiple linear regression. For each FA/MD feature in each harmonized data, we built a regression for each feature by using the same set of covariates (age, age^2^, gender, diagnosis, age*×*gender and age*×*diagnosis) as the harmonization step. We then computed *t* statistics of the estimated coefficients across all covariates and features. The boxplots of *t* statistics are shown in Figure 7. We observed that, for FA data, the magnitude of *t* statistics of all harmonized data appeared to be similar, which confirms that RELIEF did not lose biological information compared with other methods. However, for MD data, RELIEF clearly showed more significant associations with diagnosis and age*×* diagnosis than other methods, which suggests that RELIEF not only provided a thorough removal of scanner effects but also maintained biological associations well.

**Figure 7:**
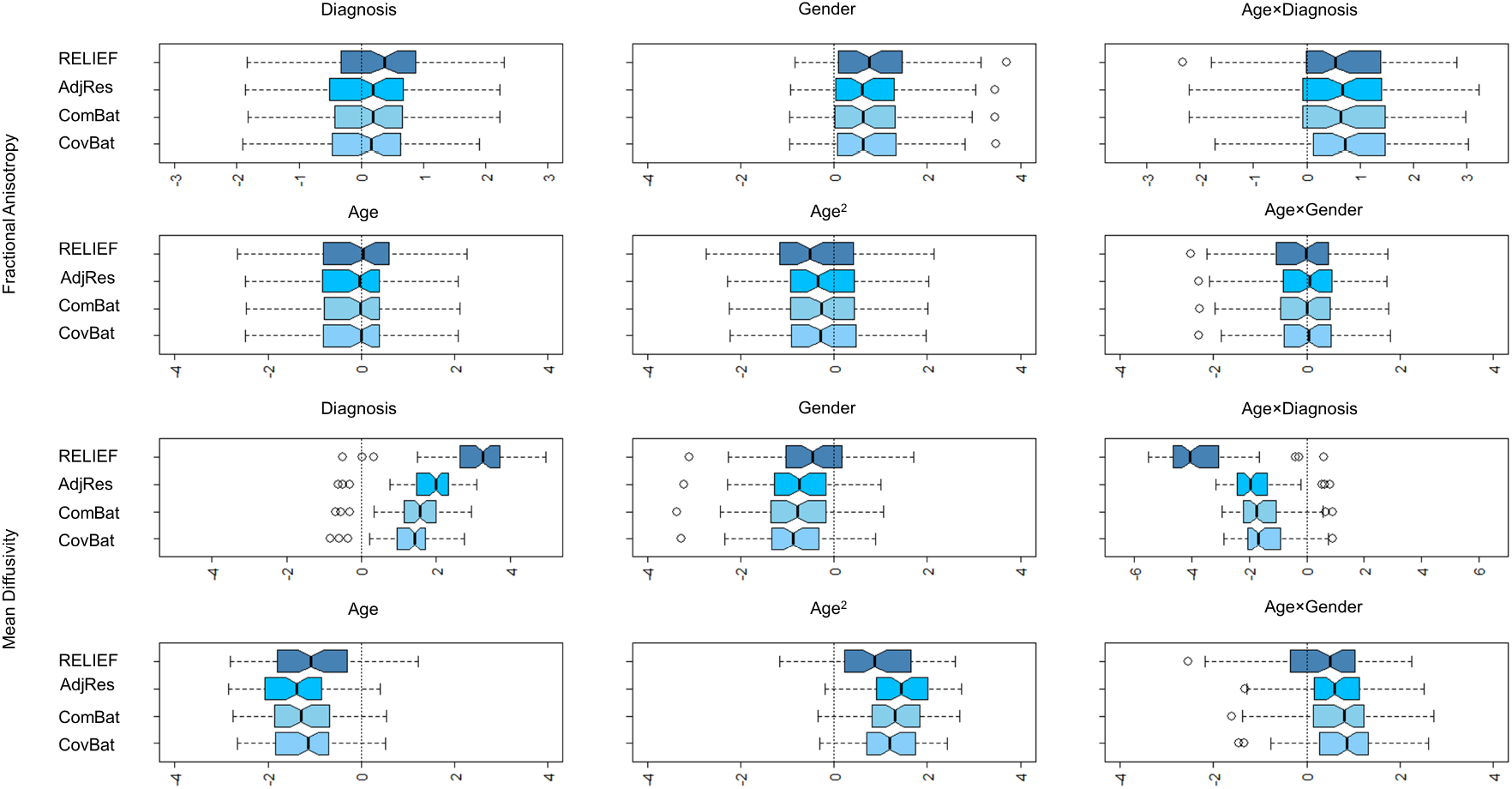
The boxplots for *t* statistics for each biological covariates used in our analysis.

## 4. Simulation Studies

### 4.1. Simulation designs

In this section, we performed extensive simulation studies to evaluate the performance of RELIEF and to compare it to other methods in controlled settings. We included ComBat, CovBat, and AdjRes as our competitors and evaluated how well-harmonized data preserves biological variations through power analysis. To evaluate the control of false positives and power, we used two models to generate heterogeneous covariances across scanners.

#### 4.1.1. Simulation 1: RELIEF model

We generated data using the sum of low-rank features following Equation (4). We simulated 1,000 null data sets with *n*_1_ = *n*_2_ = 50 (so that *n* = 100), and *V* = 100 features. Our data-generating model is summarized by

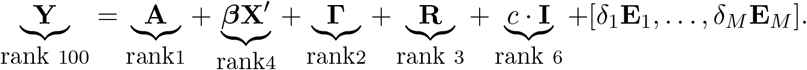

We used 4 nuisance covariates for the covariate effects, where each element of ***β*** and **A** was generated from 𝒩 (0, 1^2^). The covariate vector for each subject was generated from the multivariate normal distribution with zero means, and we used AR(1) for the covariance matrix with the autocorrelation parameter 0.2. Second, we generated **R** by first generating a *V × n* matrix whose entries are drawn from 𝒩 (0, 1^2^), then taking the first 3 principal components. Similarly, we generated each **I**_*i*_ by generating a *V × n*_*i*_ matrix using 𝒩 (0, 1^2^) then taking the top 3 principal components. Lastly, we also generated the additive scanner effect (location) *γ*_*iv*_ by fixing it to be the same for all *i* and from 𝒩 (0, 1.5^2^), and multiplicative scanner effect (scale) *δ*_*i*_ from Uniform(1, 1.5). Finally, the elements of **E** were generated from 𝒩 (0, 1^2^).

The constant *c* was chosen between 0, 1, 2, 3 to evaluate the impact of scanner-specific latent patterns on statistical power. Note that we also considered *c* = 0 to investigate whether it has comparable performance when the data-generating model does not include latent scanner effects.

#### 4.1.2. Simulation 2: CovBat model

We generated data by modifying the simulation design introduced by Chen et al. [14]. To address potential covariance scanner effects, CovBat model uses principal component (PC) scores to shift each within-scanner covariance to the pooled covariance structure. Therefore, the design aimed to evaluate whether harmonization methods can approximate the underlying covariance structure when covariance scanner effects are captured by its PC shifts.

We simulated 1,000 null data sets based on SPINS data so that *n*_1_ = 172, *n*_2_ = 179 (so that *n* = 351) and *V* = 72 features. The data *y*_*ijv*_ was generated by *y*_*ijv*_ = *α*_*v*_ + *γ*_*iv*_ + *δ*_*iv*_*ϵ*_*ijv*_, where ***α*** = (*α*_1_, …, *α*_*V*_)^*′*^ is the sample mean vector of Scanner General Electric observations in the SPINS data. The additive scanner effects ***γ***_*i*_ = (*γ*_*i*1_, …, *γ*_*iV*_)^*′*^’s are vectors drawn from 𝒩 (0, 0.1^2^). For multiplicative scanner effects, we used *δ*_1*v*_ ∼ *ℐ 𝒢* (46, 50) and *δ*_2*v*_ ∼ *I ℐ 𝒢* (51, 50) following Chen et al. [14]. From the sample correlation matrix of DTI-FA observations in the SPINS data (termed **S**) with its corresponding eigen decomposition 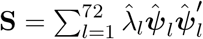, we generated ***ϵ***_*ij*_ = (*ϵ*_*ij*1_, …, *ϵ*_*ijV*_)^*′*^ that contained scanner-specific shifts. The design was to investigate how the rank of the covariance effect influences harmonization results, and we generated error terms by 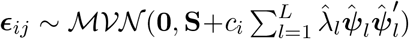, where 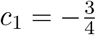 and 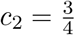. We considered different *L* including *L* = 0, 10, 20, 30.

In both simulation designs, we generated our covariate of interest, *Z*_*k*_ (*k* = 1, …, *n*), randomly from 0 or 1, for evaluation of power. We randomly chose 20% (for Simulation 1) and 50% (for Simulation 2) of features and added *τ*_*v*_ *· Z*_*k*_ to the null data, where *τ*_*v*_ ≥ 0 is the effect size for the *v*th feature, which controls whether the simulated data follows the null hypothesis *H*_0_ : *τ*_1_ = *· · ·* = *τ*_*V*_ = 0 or the alternative hypothesis *H*_1_ : at least one of *τ*_*v*_*≠* 0(*v* = 1, …, *V*). We used permutation to control family-wise error rate (FWER) at 5%.

### 4.2. Simulation results

The results for Simulation 1 are summarized in Figure 8(i). RELIEF controlled family-wise error properly, with empirical FWER of 0.044, 0.047, 0.048, and 0.052 regardless of the choice of *c*. In our simulations, while other methods controlled FWER appropriately in most scenarios, CovBat was conservative in controlling false positives when the proportion of individual latent patterns increased. In terms of power, RELIEF’s performance was nearly the same as ComBat or CovBat even when there are no latent scanner effects (i.e., *c* = 0), which supports the robustness of the proposed method. Also, as the degree of latent scanner effects (*c*) increased, RELIEF showed substantial power gain compared to others, partially because it correctly identified and removed the scanner-specific latent patterns in the data. The lower power of ComBat and AdjRes is expected as they do not consider these latent patterns in their model, and the lower power of CovBat is also expected because RELIEF’s data-generating model is different from CovBat’s assumption on PC shifts.

**Figure 8:**
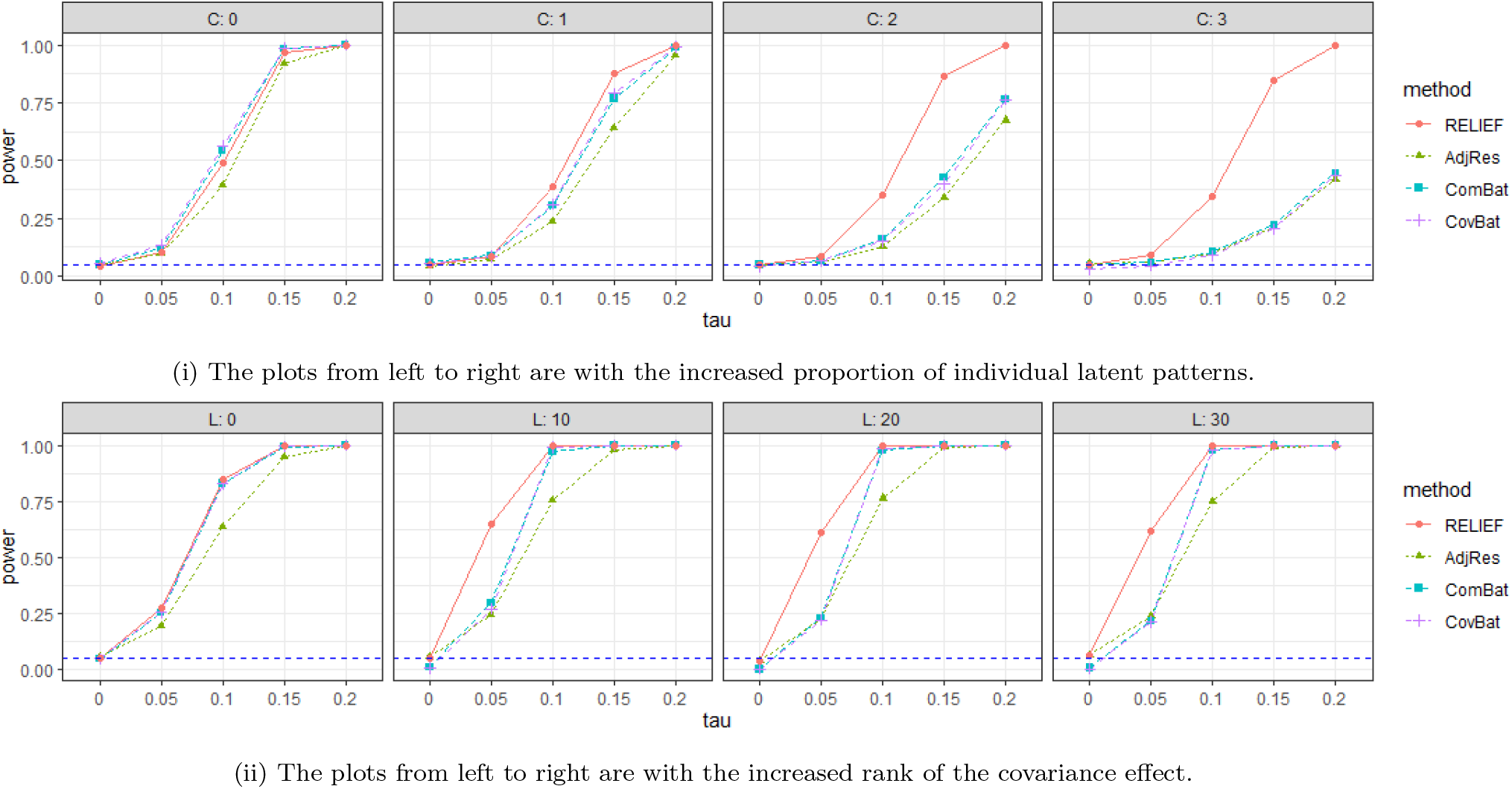
Summary of power for six harmonization methods. The blue dashed horizontal line is FWER=0.05. RELIEF controls for false positives accurately and shows superior power to competitors in both settings.

The results for Simulation 2 are summarized in Figure 8(ii). RELIEF’s empirical FWERs are 0.05, 0.051, 0.038, 0.052 for *L* = 0, 10, 20, 30, while ComBat, CovBat are conservative in controlling false positives when covariance scanner effects exist. For power, we note that when covariance scanner effects do not exist (*L* = 0), all harmonization methods increased statistical power and performed similarly, except for AdjRes whose power was lower. When *L* was large, RELIEF still showed superior performance to other methods, which supports the robustness of RELIEF even when the data-generating model did not follow the assumption of RELIEF. In addition, when SNR is low (i.e., *τ* = 0.05), RELIEF gained higher power than competitors, which supports its ability to denoise scanner effects and preserve true biological associations.

## 5. Discussion

We proposed a novel harmonization method, called RELIEF, that estimates and removes both explicit (additive and multiplicative) and latent scanner effects. RELIEF aligns with ongoing efforts to integrate neuroimaging data collected from different scanners or sites. In particular, our methods address covariance heterogeneity across different scanners, which has been a promising direction in mitigating inter-scanner biases. Our approach provides an interpretable way to harmonize heterogeneous covariances by modelling scanner-specific latent patterns under the low-rank assumption. We characterized inter-scanner bias with (i) scanner-specific means (locations), (ii) scanner-specific variances (scales), and (iii) scanner-specific latent patterns. We showed that identification of (iii), which has been overlooked in previous methods, is critical in homogenizing data from multi-site, multi-scanner neuroimaging studies.

RELIEF is a general multivariate approach that does not impose data-specific assumptions. It also does not require travelling subjects or matched controls that are often needed in supervised harmonization methods, which are infeasible in many imaging studies. Also, as we extend a regression-based approach, preserving clinical covariate effects is straightforward. Moreover, it also preserves shared variations from unobserved covariates or non-linear covariate effects using a low-dimensional representation of such variations, in which existing regression-based harmonization methods are limited.

In the analysis of the fractional anisotropy (FA) and mean diffusivity (MD) data from the SPINS study, where study samples were scanned using GE or Siemens Prisma scanners, we showed that there are substantial variations specific to Siemens Prisma. Notably, our data analysis reveals that these latent scanner effects for Siemens Prisma are heterogeneous across features (Figure 2). This result aligns with previous studies showing that inter-site variability in fractional anisotropy is specific to tissues or regions [1, 8]. RELIEF, which removed these variations in addition to the scanner-specific means and variance, successfully impaired the detection of scanners with a machine-learning method, resulting in a more homogeneous covariance as expected. A correlation analysis with existing non-biological information helped us understand the mechanism that induces these latent scanner effects.

In our simulation studies, we used different data-generating models to induce covariance heterogeneity and compared the harmonization performance of RELIEF to existing approaches. Regardless of the data-generating model used in this paper, RELIEF was superior to ComBat, CovBat, and AdjRes in preserving high statistical power. The false positive rate was also evaluated by family-wise error rate (FWER), in which RELIEF controlled it well.

RELIEF is not without limitations. First, our current approach is evaluated with a moderate number of samples. RELIEF assumes that the original data matrix consists of low-rank signals (including latent scanner effects) plus full-rank noises to scale data and choose tuning parameters. To detect these low-rank variations well, it requires a moderate number of samples to ensure the objective function of RELIEF performs more promisingly than simplified methods (e.g. ComBat) with fewer assumptions. Second, although low-rank decomposition is a useful way to capture arbitrary covariance structures, it might not always be the case when there is structured covariance in imaging data. For example, vertex-level cortical thickness data has at most 160,000 features in each brain hemisphere in FreeSurfer and reveals a high degree of spatial autocorrelation. In such a case, the low-rank assumption made in RELIEF should be evaluated carefully, and harmonizing raw images could be advantageous in accounting for anatomical variations directly [35, 36]. Also, although RELIEF does not require intense cross-validation to choose tuning parameters or ranks, it requires applying singular value decomposition (SVD) iteratively, and the computational cost increases non-linearly with increased sample size (*n*) or features (*V*). Therefore, it takes more time than existing methods (e.g. ComBat), whose computation time increase linearly with *V*. However, the computation time for RELIEF is still moderate in most downstream neuroimaging data analyses with, at most, up to hundreds of features. More importantly, we believe the powerful performance of RELIEF outweighs the cost of some additional computation time.

Also, there were recent investigations showing how pre-processing can affect the performance of ComBat harmonization, which could also be the case in RELIEF. Cetin-Karayumak et al. [24] evaluated the effect of minor differences in pre-processing on ComBat’s performance for harmonization of fractional anisotropy (FA) data across sites and showed that minor differences in the preprocessing steps resulted in non-linear changes in the input data. Because the SPINS study performed consistent preprocessing pipelines across sites, we expect its impact on our analysis to be marginal. Still, evaluating the robustness of RELIEF with respect to different pre-processing pipelines would be an interesting area of research, which we leave as future work.

RELIEF is the first approach that adopted the structured factorization of interlinked matrices into the data harmonization context, which used the concept of latent variables to characterize scanner effects. In the past decade, there have been a number of methodological developments in linked matrix factorization [20, 37, 21], which provided novel insights into understanding multimodal data [23], disease subtypes, or clustering. We believe more methodological research on data harmonization from the viewpoint of the linked matrix factorization would lead to further improvements in the harmonization quality.

To summarize, we proposed a new harmonization method, RELIEF, that contributes to ongoing efforts on integrating heterogeneous multi-site, multi-scanner studies in neuroimaging. Our novel contribution is the development of a multivariate harmonization method that captures scanner-specific latent factors, which have not been addressed in existing methods. With the three-source characterization of inter-scanner biases (location, scale, latent), RELIEF shows promising results in harmonizing all of them, eventually resulting in higher power in association studies than existing harmonization methods.

## 6. Software

RELIEF is made publicly available as an R package on GitHub: https://github.com/junjypark/RELIEF. It requires the same input as neuroComBat [38] (imaging data matrix, covariates, and scanner information), producing harmonized imaging data in the same format. Our harmonization took approximately 4 seconds on a Macbook Pro 2018 to harmonize data with 72 imaging features from 351 subjects, which supports the computational efficiency of the proposed method.

## Acknowledgements

The authors would like to thank all participants for their contribution to this work, and the research staff who performed data collection and management. The SPINS study was supported by the National Institute of Mental Health (1/3R01MH102324-01, 2/3R01MH102313-01, 3/3R01MH102318-01). LDO was supported by the Brain & Behavior Research Foundation. ANV was supported by the National Institute of Mental Health (1/3R01MH102324 & 1/5R01MH114970), Canadian Institutes of Health Research, Canada Foundation for Innovation, CAMH Foundation, and University of Toronto. JYP was supported by Natural Sciences and Engineering Research Council of Canada (NSERC) (RGPIN-2022-04831) and the University of Toronto’s Data Science Institute (Catalyst Grant).

## References

[1] C. Vollmar, J. O’Muircheartaigh, G. J. Barker, M. R. Symms, P. Thompson, V. Kumari, J. S. Duncan, M. P. Richardson, M. J. Koepp, Identical, but not the same: intra-site and inter-site reproducibility of fractional anisotropy measures on two 3.0 T scanners, Neuroimage 51 (2010) 1384–1394.

[2] T. Zhu, R. Hu, X. Qiu, M. Taylor, Y. Tso, C. Yiannoutsos, B. Navia, S. Mori, S. Ekholm, G. Schifitto, et al., Quantification of accuracy and precision of multi-center DTI measurements: a diffusion phantom and human brain study, Neuroimage 56 (2011) 1398–1411.

[3] X. Han, J. Jovicich, D. Salat, A. van der Kouwe, B. Quinn, S. Czanner, E. Busa, J. Pacheco, M. Albert, R. Killiany, et al., Reliability of MRI-derived measurements of human cerebral cortical thickness: the effects of field strength, scanner upgrade and manufacturer, Neuroimage 32 (2006) 180–194.

[4] H. Takao, N. Hayashi, K. Ohtomo, Effects of study design in multi-scanner voxel-based morphometry studies, Neuroimage 84 (2014) 133–140.

[5] C. Dansereau, Y. Benhajali, C. Risterucci, E. M. Pich, P. Orban, D. Arnold, P. Bellec, Statistical power and prediction accuracy in multisite resting-state fMRI connectivity, Neuroimage 149 (2017) 220–232.

[6] W. E. Johnson, C. Li, A. Rabinovic, Adjusting batch effects in microarray expression data using empirical Bayes methods, Biostatistics 8 (2007) 118–127.

[7] Y. Zhang, G. Parmigiani, W. E. Johnson, ComBat-seq: batch effect adjustment for RNA-seq count data, NAR Genomics and Bioinformatics 2 (2020) qaa078.

[8] J.-P. Fortin, D. Parker, B. Tunç, T. Watanabe, M. A. Elliott, K. Ruparel, D. R. Roalf, T. D. Satterthwaite, R. C. Gur, R. E. Gur, et al., Harmonization of multi-site diffusion tensor imaging data, NeuroImage 161 (2017) 149–170.

[9] J.-P. Fortin, N. Cullen, Y. I. Sheline, W. D. Taylor, I. Aselcioglu, P. A. Cook, P. Adams, C. Cooper, M. Fava, P. J. McGrath, et al., Harmonization of cortical thickness measurements across scanners and sites, NeuroImage 167 (2018) 104–120.

[10] M. Yu, K. A. Linn, P. A. Cook, M. L. Phillips, M. McInnis, M. Fava, M. H. Trivedi, M. M. Weissman, R. T. Shinohara, Y. I. Sheline, Statistical harmonization corrects site effects in functional connectivity measurements from multi-site fMRI data, Human Brain Mapping 39 (2018) 4213–4227.

[11] J. C. Beer, N. J. Tustison, P. A. Cook, C. Davatzikos, Y. I. Sheline, R. T. Shinohara, K. A. Linn, A. D. N. Initiative, et al., Longitudinal ComBat: A method for harmonizing longitudinal multi-scanner imaging data, NeuroImage 220 (2020) 117129.

[12] R. Pomponio, G. Erus, M. Habes, J. Doshi, D. Srinivasan, E. Mamourian, V. Bashyam, I. M. Nasrallah, T. D. Satterthwaite, Y. Fan, et al., Harmonization of large MRI datasets for the analysis of brain imaging patterns throughout the lifespan, NeuroImage 208 (2020) 116450.

[13] A. A. Chen, C. Luo, Y. Chen, R. T. Shinohara, H. Shou, A. D. N. Initiative, et al., Privacy-preserving harmonization via distributed ComBat, NeuroImage 248 (2022) 118822.

[14] A. A. Chen, J. C. Beer, N. J. Tustison, P. A. Cook, R. T. Shinohara, H. Shou, A. D. N. Initiative, Mitigating site effects in covariance for machine learning in neuroimaging data, Human Brain Mapping 43 (2022) 1179–1195.

[15] A. A. Chen, D. Srinivasan, R. Pomponio, Y. Fan, I. M. Nasrallah, S. M. Resnick, L. L. Beason-Held, C. Davatzikos, T. D. Satterthwaite, D. S. Bassett, et al., Harmonizing functional connectivity reduces scanner effects in community detection, NeuroImage 256 (2022) 119198.

[16] J. T. Leek, J. D. Storey, Capturing heterogeneity in gene expression studies by surrogate variable analysis, PLoS Genetics 3 (2007) e161.

[17] J.-P. Fortin, E. M. Sweeney, J. Muschelli, C. M. Crainiceanu, R. T. Shinohara, A. D. N. Initiative, et al., Removing inter-subject technical variability in magnetic resonance imaging studies, NeuroImage 132 (2016) 198–212.

[18] J. A. Gagnon-Bartsch, L. Jacob, T. P. Speed, Removing unwanted variation from high dimensional data with negative controls, Berkeley: Tech Reports from Dep Stat Univ California (2013) 1–112.

[19] J. Y. Park, E. F. Lock, Integrative factorization of bidimensionally linked matrices, Biometrics 76 (2020) 61–74.

[20] E. F. Lock, K. A. Hoadley, J. S. Marron, A. B. Nobel, Joint and individual variation explained (JIVE) for integrated analysis of multiple data types, The Annals of Applied Statistics 7 (2013) 523.

[21] I. Gaynanova, G. Li, Structural learning and integrative decomposition of multi-view data, Biometrics 75 (2019) 1121–1132.

[22] E. F. Lock, J. Y. Park, K. A. Hoadley, Bidimensional linked matrix factorization for pan-omics pan-cancer analysis, The Annals of Applied Statistics 16 (2022) 193.

[23] Q. Yu, B. B. Risk, K. Zhang, J. Marron, JIVE integration of imaging and behavioral data, NeuroImage 152 (2017) 38–49.

[24] S. Cetin-Karayumak, K. Stegmayer, S. Walther, P. R. Szeszko, T. Crow, A. James, M. Keshavan, M. Kubicki, Y. Rathi, Exploring the limits of ComBat method for multi-site diffusion MRI harmonization, bioRxiv (2020) 2020–11.

[25] M. Gavish, D. L. Donoho, Optimal shrinkage of singular values, IEEE Transactions on Information Theory 63 (2017) 2137–2152.

[26] T. Hastie, R. Mazumder, J. D. Lee, R. Zadeh, Matrix completion and low-rank SVD via fast alternating least squares, The Journal of Machine Learning Research 16 (2015) 3367–3402.

[27] V. Nygaard, E. A. Rødland, E. Hovig, Methods that remove batch effects while retaining group differences may lead to exaggerated confidence in downstream analyses, Biostatistics 17 (2016) 29–39.

[28] J.-D. Tournier, R. Smith, D. Raffelt, R. Tabbara, T. Dhollander, M. Pietsch, D. Christiaens, B. Jeurissen, C.-H. Yeh, A. Connelly, Mrtrix3: A fast, flexible and open software framework for medical image processing and visualisation, Neuroimage 202 (2019) 116137.

[29] C. Bhushan, J. P. Haldar, S. Choi, A. A. Joshi, D. W. Shattuck, R. M. Leahy, Co-registration and distortion correction of diffusion and anatomical images based on inverse contrast normalization, Neuroimage 115 (2015) 269–280.

[30] J. G. Malcolm, M. E. Shenton, Y. Rathi, Filtered multitensor tractography, IEEE transactions on medical imaging 29 (2010) 1664–1675.

[31] F. Zhang, Y. Wu, I. Norton, L. Rigolo, Y. Rathi, N. Makris, L. J. O’Donnell, An anatomically curated fiber clustering white matter atlas for consistent white matter tract parcellation across the lifespan, NeuroImage 179 (2018) 429–447.

[32] F. Zhang, Y. Wu, I. Norton, Y. Rathi, A. J. Golby, L. J. O’Donnell, Test–retest reproducibility of white matter parcellation using diffusion mri tractography fiber clustering, Human brain mapping 40 (2019) 3041–3057.

[33] C. Beaulieu, The basis of anisotropic water diffusion in the nervous system–a technical review, NMR in Biomedicine: An International Journal Devoted to the Development and Application of Magnetic Resonance In Vivo 15 (2002) 435–455.

[34] L. J. O’Donnell, C.-F. Westin, An introduction to diffusion tensor image analysis, Neurosurgery Clinics 22 (2011) 185–196.

[35] H. Mirzaalian, L. Ning, P. Savadjiev, O. Pasternak, S. Bouix, O. Michailovich, G. Grant, C. E. Marx, R. A. Morey, L. A. Flashman, et al., Inter-site and inter-scanner diffusion MRI data harmonization, NeuroImage 135 (2016) 311–323.

[36] S. C. Karayumak, S. Bouix, L. Ning, A. James, T. Crow, M. Shenton, M. Kubicki, Y. Rathi, Retrospective harmonization of multi-site diffusion mri data acquired with different acquisition parameters, Neuroimage 184 (2019) 180–200.

[37] Q. Feng, M. Jiang, J. Hannig, J. Marron, Angle-based joint and individual variation explained, Journal of Multivariate Analysis 166 (2018) 241–265.

[38] J. Fortin, Harmonization of Multi-Site Imaging Data with ComBat, R Package Version 1.0. 9. neuroCombat., 2021.

